# An updated population estimate for Northern gannets across their north-east Atlantic breeding range following the 2022 outbreak of High pathogenicity avian influenza

**DOI:** 10.64898/2026.04.24.720534

**Authors:** L. Quinn, J.W.E. Jeglinski, S Auhage, D. Balmer, I.S. Bringsvor, E. Burton, J.H.F. Castenschiold, S. Christensen-Dalsgaard, J. Danielsen, J. Dierschke, A.V. Ezhov, G. A. Guðmundsson, T. Hart, M. Jessopp, R. Jones, Y.V. Krasnov, S-H. Lorentsen, A. Palsdottir, P. Provost, A. Purdie, G.D. Morgan, E. Murphy, M. Murphy, B. Olsen, E. Owen, H. Strøm, I. Støyle Bringsvor, T.D. Tierney, L.J. Wilson, S. Wanless

**Affiliations:** NatureScot, Great Glen House, Inverness, UK; Institute of Ecoscience, Aarhus University, Denmark; School of Biodiversity, One Health and Veterinary Medicine, University of Glasgow, UK; British Trust for Ornithology, The Nunnery, Thetford; Scottish Seabird Centre, North Berwick; Norwegian Institute for Nature Research NINA, Trondheim, Norway; Faeroe Marine Research Institute Havstovan, Tórshavn, Faroe Islands; Institut fuer Vogelforschung ‘Vogelwarte Helgoland‘, Helgoland, Germany; Murmansk Marine Biological Institute of the Russian Academy of Sciences, Russia; Natural Science Institute of Iceland, Gardabaer, Iceland; School of Biomedical Sciences, Oxford Brookes University, Headington Campus, Oxford, OX3 0BP; School of Biological, Earth & Environmental Sciences, University College Cork, Ireland; Natural England, Foss House, YO1 7PX; Ligue pour la Protection des Oiseaux, Réserve Naturelle Nationale des Sept-Iles, Pleumeur Bodou, France; Alderney Wildlife Trust, St. Anne, Alderney, Bailiwick of Guernsey; Sustainability Institute, University College Cork, Ireland; Natural Resources Wales, Maes y Ffynnon, Penrhosgarnedd, Bangor, Gwynedd, LL57 2DW; The National Trust for Scotland, Balnain House, Huntly Street Inverness IV3 5HR; BirdLife Møre og Romsdal, Sandshamn, Norway; Norwegian Polar Institute, Post box 6606 Stakkevollan, Tromsø, Norway; National Parks and Wildlife Service, Dublin, Ireland; RSPB Centre for Conservation Science, Etive House, Inverness, IV2 3BW, UK; UK Centre for Ecology & Hydrology, Bush Estate, Penicuik, UK, EH26 0QB; RSPB Ramsey Island, St Davids, Pembrokeshire, SA62 6PY; Kandalaksha State Nature Reserve, Russia

**Keywords:** Northern gannet, HPAI, census, metapopulation

## Abstract

Northern gannets (*Morus bassanus*) have been regarded as a seabird ‘success’ story, due to population increases throughout the 20^th^ and 21^st^ century contrasting with global seabird declines. However, in 2022 gannets experienced a severe outbreak of High Pathogenicity Avian Influenza (HPAI) across their global distribution, leading to an urgent need to reassess their population status. This study presents breeding gannet census numbers for 2023/24 from all colonies across the North-East Atlantic metapopulation (Great Britain, Ireland, the Channel Islands, Iceland, Norway, the Faroe Islands, France, Germany, Russia). Gannet numbers decreased by 17% across the North-East Atlantic metapopulation between the 2013/14 and 2023/24 census from 414,598 to 345,854 apparently occupied sites (AOS), a global decrease of at least 13%. The bulk of the reduction in AOS was driven by the largest colonies (>10,000 AOS) each losing tens of thousands of AOS. These figures likely underestimate the impact of the HPAI outbreak worldwide, since most colonies will have increased between the last census in 2013/14 and the 2022 HPAI outbreak, and the Canadian breeding population was last counted pre-HPAI outbreak. Scotland still holds the largest proportion of both the North-East Atlantic metapopulation (59%), and the world population (46%), while Great Britain, Ireland and the Channel Islands together hold 83% of the North-East Atlantic metapopulation and 64% of the world population. This study not only presents an updated population census for gannets in the North-East Atlantic but illustrates the large-scale impacts of a disease outbreak on a seabird species across its global range and highlights the importance of more regular census efforts to better quantify the demographic consequences of such events.

## Introduction

Northern gannet (*Morus bassanus*, hereafter ‘gannet’) colonies in Great Britain (GB), Ireland and the Channel Islands form the majority of the North-East Atlantic metapopulation, which extends geographically from northern France and northern Germany in the south, to Iceland, northern Norway, Murman coast (Russia) and Svalbard in the north (Jeglinski *et al*. 2023; 2024). The North-East Atlantic metapopulation consisted of 46 colonies in 2016 (Jeglinski *et al*. 2023) and is considered distinct from the North-West Atlantic metapopulation, which is made up of six Canadian colonies (Nelson 2002; Wanless *et al*. 2023). Previous censuses estimated that GB, Ireland and the Channel Island colonies held around 84% and 66% of the North-East Atlantic metapopulation and global population, respectively (Murray *et al*. 2015). Due to the high proportion of their global breeding population occurring in the GB and the Channel Islands region, gannets are amber on the Birds of Conservation Concern (BoCC) list (Stanbury *et al*. 2024). Scotland holds not only the oldest mentioned but also the world’s largest gannet colonies (Gurney 1913; Wanless *et al*. 2023) while the colonies at the southern and northern fringes of the metapopulation are smaller and provide additional important evidence of ancient colonisations (Iceland and Faroe Islands), recent range expansions (Germany, France, Norway and Russia), and, compared to the rest of the metapopulation, highly unusual colonisation-extinction dynamics (Norway) (Barrett 2008; Barrett *et al*. 2017; Jeglinski *et al*. 2024).

Gannets have been censused approximately every ten years across GB and Ireland since the 1980s (Murray *et al*. 2015), though much longer time series exist for many of the older colonies (Gurney 1913, Fisher & Vevers 1943, Nelson 2002). The last gannet census in 2013/14 showed an increase in gannet populations across their metapopulation as a whole (Murray *et al*. 2015; Wanless *et al*. 2023). However, throughout their global distribution, gannets were significantly impacted by a severe outbreak of High Pathogenicity Avian Influenza (hereafter HPAI) (Lane *et al*. 2024) in 2022. Evidence of population declines was apparent in a subset of colonies surveyed in 2022 and 2023 (Tremlett *et al*. 2024; Atkinson *et al*. 2025; Ezhov & Krasnov 2024). An updated census across all remaining GB, Ireland and the Channel Islands, as well as across the wider North-East Atlantic populations is crucial to assess the wider impact of HPAI in gannet colonies.

Here, we present the results of the latest GB, Ireland and the Channel Islands gannet census, which was undertaken during the 2023 and 2024 breeding seasons, following the 2022 HPAI outbreak, and compare these with the results of previous censuses. We set these findings in the context of the wider North-East Atlantic metapopulation by collating the most recent available data on colony sizes and colonisation attempts across the rest of their North-East Atlantic range, including Iceland, Norway, the Faroe Islands, France, Russia, and Germany. We also compare population trends across their range, where sufficient time series data are available (1995 onwards). Finally, we provide a revised global estimate although we caution to treat this as an overestimate since no updated counts for the Canadian population have been published following the 2022 HPAI outbreak.

## Methods

### Census survey years

We based our analysis on census data from 2023 wherever possible (43 of 50 colonies). Twelve colonies were counted in both 2023 and 2024 (Supplementary Table S3). Where data in 2023 were deemed insufficient in quality, we were therefore able to use census data from 2024, e.g. for Ailsa Craig and Sula Sgeir we selected 2024 instead of 2023 as image quality was higher and therefore inter-observer variation in counts was lower. If no surveys had taken place in either year, we used the latest available count instead, resulting in one Icelandic and four Norwegian colonies having a count either before or after 2023/24 (see Supplementary Table S2).

### Survey methods

All known gannet colonies across the North-East Atlantic range were surveyed (n=50, Figure 1), using a combination of land, boat, aircraft or drone (see Supplementary Table S1 for full summary of survey methods).

**Figure 1:**
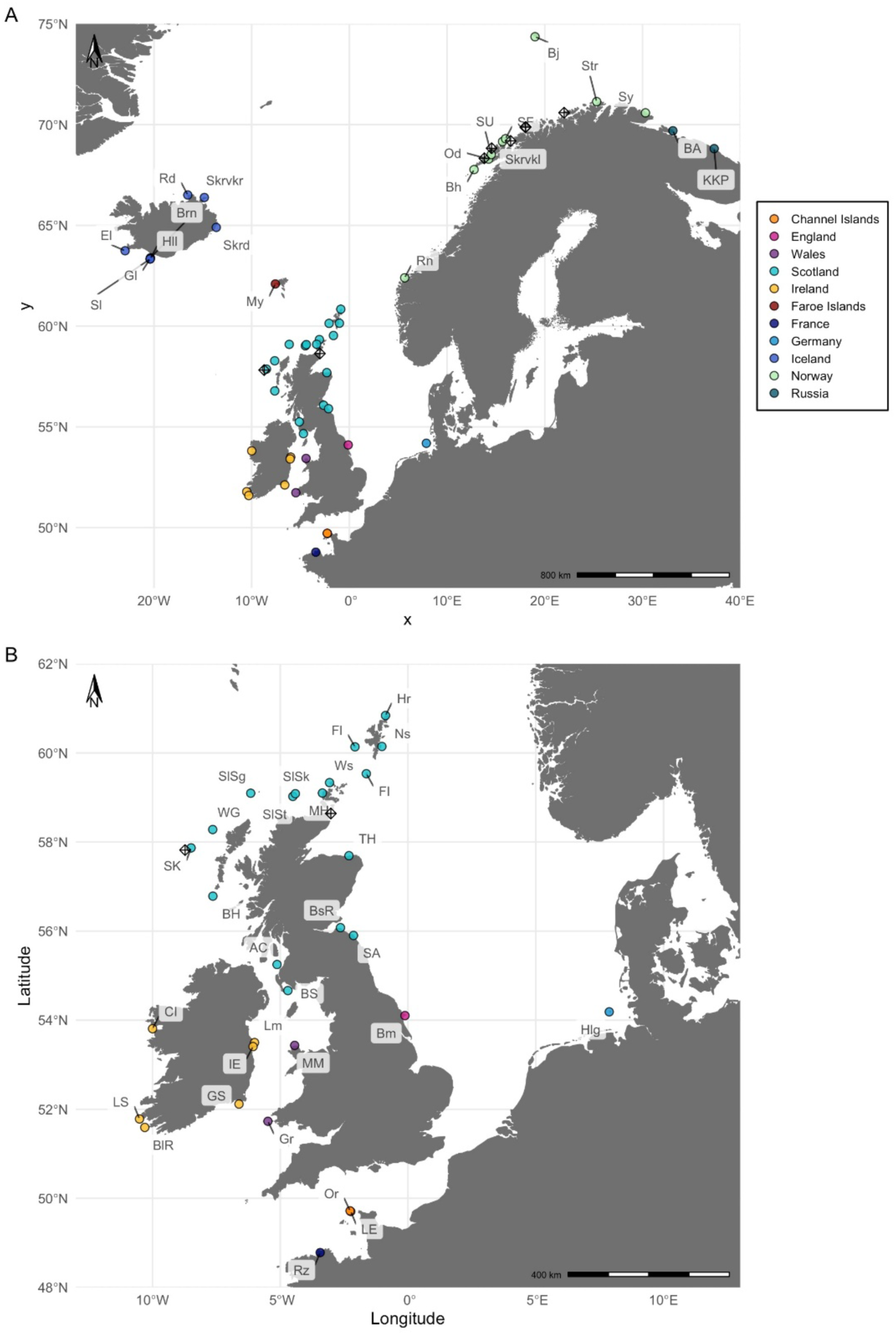
A) Map of all known current gannet colonies in the North-East Atlantic metapopulation with B) inset of the southern central region. Colonies are indicated with dots coloured by geographic region (see legend), colonisation attempts (for definition see Methods) are indicated with black diamonds. For full spelling of abbreviations see Supplementary Table S2.

The data for GB, Ireland and the Channel Islands colonies collected as part of this study were extracted from the Seabird Monitoring Programme database (Accessed 21^st^ April 2025). Reports and papers produced on specific colonies go into further methodological detail, references for which are provided within Supplementary Table S1. Data for the other countries within the North-East Atlantic were kindly supplied by the respective researchers and institutions and collated for this paper. Detailed survey methods for the other countries within the North-East Atlantic are not presented, but the survey type is summarised in Supplementary material Table S1. Long-term data time series were taken from Jeglinski *et al*. (2023).

The census unit for most colonies was Apparently Occupied Sites (AOS), with 10 of the 50 total colonies surveyed recording Apparently Occupied Nests (AON). AOS and AON were identified based on the definition in the Seabird Monitoring Handbook. AOS are defined as *‘a site occupied by one or two adult gannets irrespective of whether or not any nesting material is present, so long as the site appears suitable for breeding. Sites with unattended chicks are included’* (Walsh *et al*. 1995). AON are defined as *‘one or two adults with nest material, or a large chick, are present’* (Walsh *et al*. 1995). We assumed AONs were equivalent to AOS, i.e. no correction factor was required, nor is it recommended (Walsh *et al*. 1995). Using AOS means there may be a slight overestimation in the counts, as it does not require nest material to be visible, unlike AONs, but it is the most appropriate unit to use, especially when surveying from a distance. Therefore, the overall gannet count presented uses AOS units.

We distinguish colonisation attempts, i.e. observations of one to two gannet AOS for one to two years, generally ephemeral, from colonies, i.e. stable breeding sites occupied by more than two gannet AOS for at least three years. Note that we follow Jeglinski *et al*. (2023) in classifying Rockall as a colonisation attempt despite a longer time series of observations of larger numbers of AOS due to the ephemeral nature of the exposed rock which makes a stable breeding colony unlikely.

#### Land or boat-based survey methods

Traditional methods using land or boat-based counts followed Walsh *et al*. 1995 and were consistent with those used during previous land and boat-based surveys (Murray *et al*. 2015). Fourteen of the 50 colonies carried out land or boat-based counts with observers in situ, and for two colonies (Hermaness and Noss) counts also included colony sections that were photographed using a DSLR camera (Sony A35 for Hermaness and Nikon D7200 for Noss) and then retrospectively counted (see Supplementary Table S1 for further detail). Software used for processing and counting images for these two colonies was Adobe Photoshop Version 25.0, 2023.

#### Fixed-wing or helicopter aerial surveys

For all aerial surveys using fixed wing or helicopters a 100% coverage was achieved. Aerial surveys were the most common survey method, being carried out for 33 of the 50 colonies. For the five western Scotland colonies (Flannan Isles, Sule Skerry, Sule Stack, Sula Sgeir, Rockall) the survey followed the same methods as the 2013 gannet census (Wanless *et al*. 2015), with an additional vertical camera (Olaya Meza & Jones 2025). During the 2023/24 census no ‘controlled disturbance’ flights were carried out to flush non-breeders, as had been carried out in 2013, because higher image quality now allows loafing birds to be identified and therefore excluded from the count. The Scottish aerial survey was carried out by a twin-engine fixed-wing aircraft that orbited the target area between 1,300 and 1,500 ft, capturing data using a handheld oblique camera (Canon 5D with a 200mm prime lens) positioned at the aircraft side window. The focal length achieved a Ground Sampling Distance (GSD) of between 1.5-2.0 centimetres (cm). Where AOS were obscured by terrain, the vertical imagery camera was checked to ensure complete coverage. All aerial surveys of Irish colonies took place using a helicopter with a Nikon D7200 camera with a 24-120mm lens operated by an observer seated in the front left position of the aircraft beside the pilot to obtain an unobscured view and advise on the best approach to limit disturbance. The pilot circled the colonies rather than flying directly over them. All aerial surveys for the Channel Islands colonies used a fixed-wing plane with the photographer seated alongside the pilot, photographing through an open window hatch using a Nikon D800 DSLR camera with a 70-200 mm lens. The pilot circled the colonies rather than flying directly over them. An example image from the aerial surveys is shown in Supplementary material Figure S1 and S2.

Software used for viewing and processing images from aerial surveys included: Geographic Information Systems (GIS) QGIS software (version 3.28, 2022), Adobe Photoshop (Version 25.0, 2023), GIMP (Version 2.10.24, 2021, https://www.gimp.org/); and for AOS annotation and subsequent count by ornithologists - DotDotGoose (version 1.6.0, 2023, biodiversityinformatics.amnh.org/open_source/dotdotgoose/).

#### Drone-based aerial survey methods

In contrast to previous censuses where a combination of traditional land-based, boat-based and aerial surveys were employed, the 2023/2024 census saw the introduction of drone-based surveys at 12 of the 50 colonies (see Supplementary Table S1). Survey methods followed guidelines within Edney *et al*. 2023. Prior to the census, discussions between drone surveyors across GB further ensured consistent methodologies across colonies.

Drone surveys involved different drone model types (Supplementary Table S1). Drone surveys within protected sites were carried out with landowner permission and relevant licences. Flights were kept to 100m distance where possible, and did not approach closer than 50m, as recommended by Edney *et al*. (2023). An additional person (‘spotter’) accompanied the drone pilot to assist in ensuring disturbance was minimised. Flights were generally flown in preprogrammed transects with at least 75% overlap between images (except for St Kilda, see Nisbet *et al*. 2015, and the Faroes was at least 65% overlap J. Castenschiold *pers comm.*). All sites where drones were used had 100% coverage, except Bass Rock where a small cliff area was not covered. In this case, an assumption that the dispersion of AOS in 2023 was the same as in 2014 was made, the count was then scaled up to give a total population estimate (further details in Harris *et al*. 2023).

Post-image analysis for drone imagery of gannet colonies used a number of different stitching and/or mapping softwares, including: Panorama stitcher v1.11, Maps Made Easy (accessed across various dates from https://www.mapsmadeeasy.com/), Drone2Map 2025.1, Agisoft Metashape (v2.1.4; Agisoft LLC, 2025), and GIS mapping softwares (including ArcGIS 11.0, ArcGIS Pro v.3, and QGIS 3.40) to create high-resolution orthomosaic images. An example of a stitched image is provided in Supplementary material Figure S3 and S4.

### Analysis

With any count data there is an element of count error associated with the survey method due to observer differences in detecting and classifying AOS. To reduce count error, only images deemed to have suitable image quality (fixed wing aerial or drone imagery) were used. Images were counted by at least two ornithologists and were deemed to be accurate if independent counts were within 10% of one another and were then averaged across counters.

We calculated the ‘percentage change’ to quantify the change in colony size (AOS) between the most recent survey (2023/24) and the last survey (2013/14) for the entire North-East Atlantic metapopulation, noting that for four Norwegian colonies the most recent surveys were prior to the 2022 HPAI outbreak. To put these estimates into a global context, we use the most recent published estimate for the North-West Atlantic metapopulation (Wanless *et al*. 2023), noting that these census data predate the 2022 HPAI outbreak. For comparison with the 2013/14 and the 2003/04 census, data presented in the detailed geographic accounts tables was taken from Murray *et al*. 2015 or from Jeglinski *et al*. 2023. We explored possible explanations for the variation in colony size changes across the North-East Atlantic metapopulation. We investigated if the percent changes or absolute changes (the difference in colony size between both census years) in colony size were driven by large-scale geographic patterns, the previous size of the colony or the first date of observing unusual mortality as an indicator of the HPAI outbreak in 2022 (Lane *et al*. 2024) by running a general linear model (glm) with percent change or absolute change as response variable, and latitude, previous colony size and date of first HPAI outbreak detection as explanatory variables. We fitted all three covariates as additive effects, and simplified and compared models using AIC (Burnham & Anderson 2004). All data processing, statistical modelling and mapping was performed in R version 4.5.2 (R Core Team, 2025). Time series plots and maps were produced using R packages *ggplot* and *ggspatial*, respectively (Wickham 2016; Dunnington 2025), the glm was run using the *stats* package in the base implementation of R (R Core team, 2025) and the land shapefile used for mapping was acquired using the R package *rnaturalearth* (Massicotte & South 2025).

## Results

### North-East Atlantic and global population estimates

The gannet population counts show a decrease of 17% across the North-East Atlantic metapopulation and of 13% across the global population from 2013/14 to 2023/24 (Table 1). This global population estimate should be interpreted with caution, as the Canadian colonies have not had a complete census since 2018-2020 (before HPAI) and thus available count data for these colonies do not include any HPAI-related decreases. Previously, Canadian colonies totalled 21% of the global population (Murray *et al*. 2015). It is therefore likely that the world population estimate stated here is an overestimation. Scotland still holds the largest proportion of the world population, with 59% of the North-East Atlantic population and 46% of the world population breeding there.

**Table 1.**
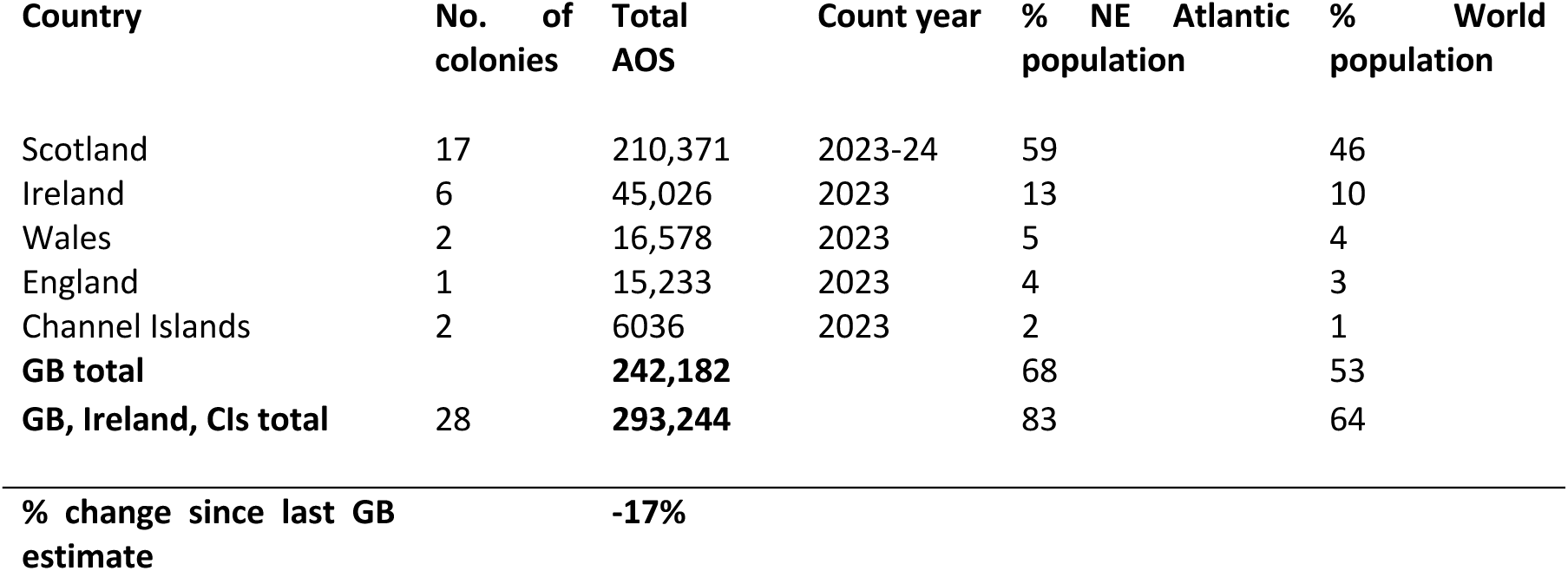
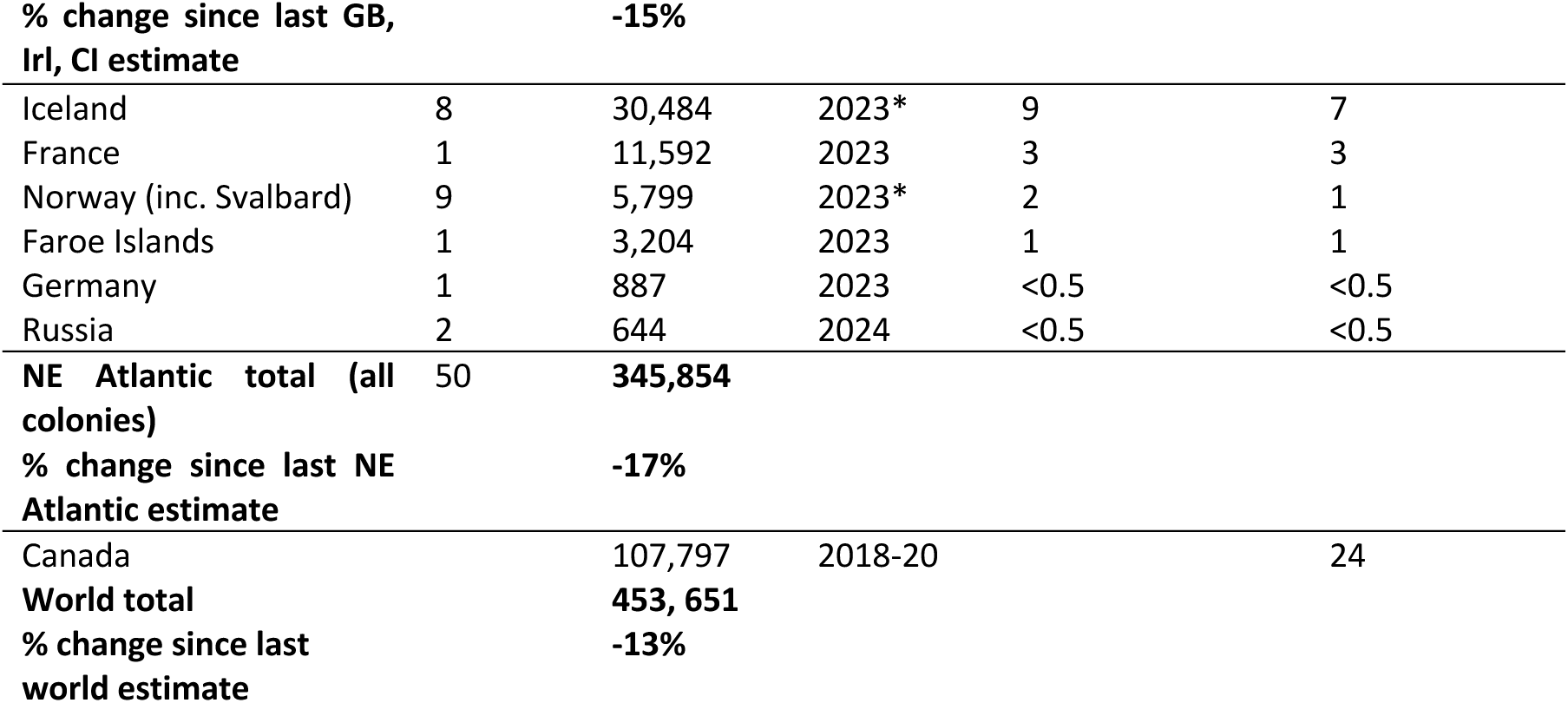
Overall population estimates of gannet. Data from all North-East (NE) Atlantic colonies have been collated or collected as part of this study. The Canadian population estimate is taken from Wanless *et al*. 2023 and is based on data collected 2018-2020. *Not all Norwegian colonies had census data in 2023, for these the latest available date was chosen, see Table S1 in supplementary material. *One Icelandic colony used a count from 2025. Percentages are rounded to the nearest whole number.

The absolute decrease in AOS in the North-East Atlantic metapopulation was driven by the largest colonies (more than 10,000 AOS, Figure 2C top) which each lost tens of thousands of AOS, particularly evident for the Bass Rock, Grassholm, Little Skellig, Ailsa Craig, Hermaness, Skrudur and Rouzic. The size of a colony in the previous census significantly drove absolute changes in AOS between both census years (glm, previous colony size, t = 8.015, p < 0.01, r^2^ = 0.584) while latitude or the date of first detection of unusual mortality in 2022 did not (glm, p > 0.05). Among the large colonies, these severe losses contrasted with sizeable increases in absolute numbers of AOS at Bempton and Skrudur (Figure 2C). Among mid-sized and smaller colonies, the direction and magnitude of changes in absolute numbers were highly variable, with no clear pattern in increases or decreases.

**Figure 2:**
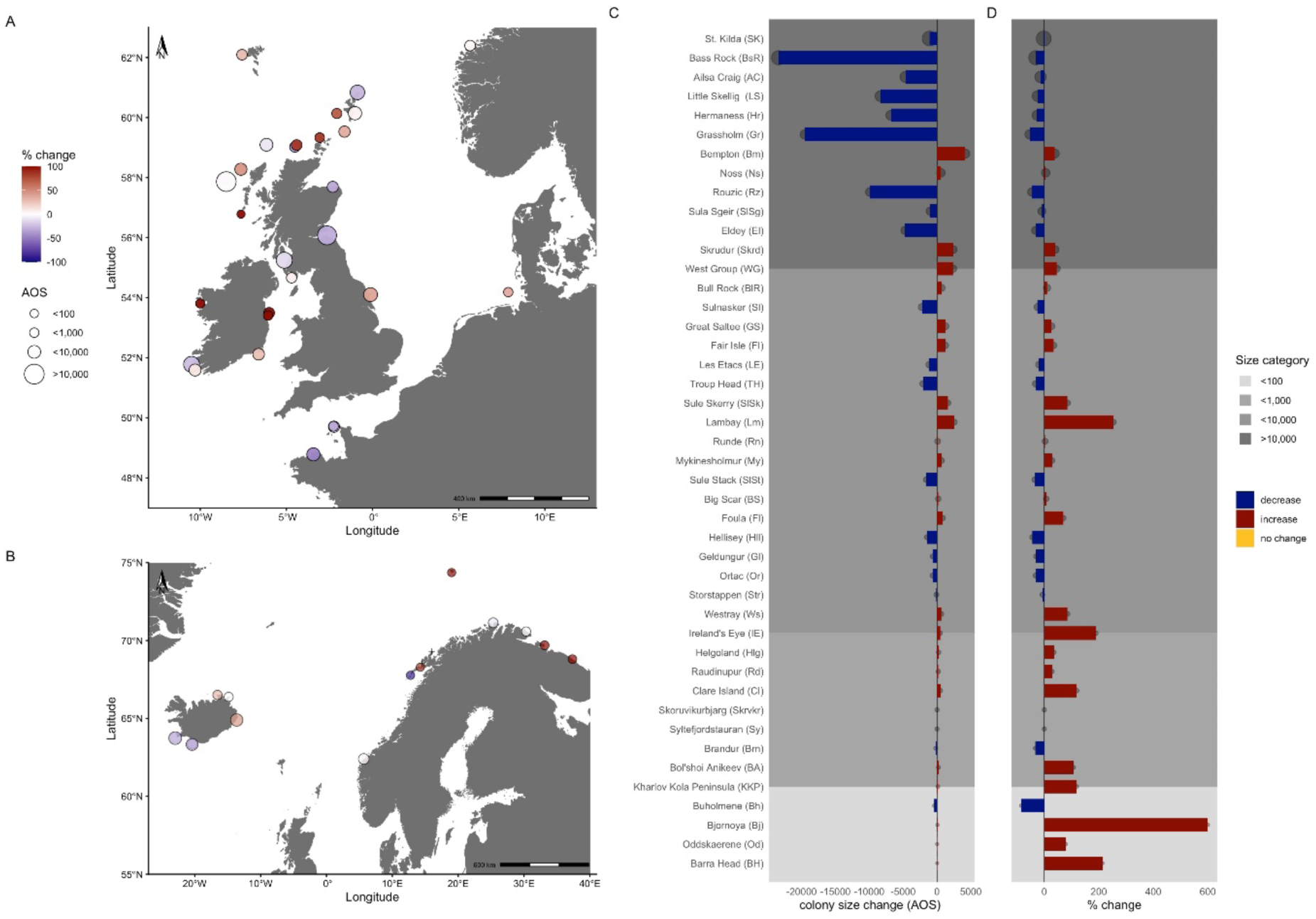
All 50 gannet colonies in the A) southern and central, B) northern part of the North-East Atlantic metapopulation. The point size illustrates the colony size (binned) at the 2023/24 census, and the colour indicates the direction and strength of change in size between 2013/14 and 2023/24 censuses), blue = colony size decrease; red = colony size increase. Note that % change is curtailed to 100% to avoid a few small colonies with large % increases (e.g. Bjørnøya 630 % change) driving the colour scale. For absolute values and complete colony names, see Supplementary Table S2). C) absolute differences in AOS and C) percent differences in colony size between the last two censuses (2013/14 and 2023/24) for all colonies in the North-East Atlantic metapopulation, for exceptions see able S2 in supplementary materials. Bar colour and orientation indicate direction of change, the size of the associated dots indicates colony size. Bars are ordered by colony size (2023/24 census), in descending order. Background colour in increasingly lighter shades of grey indicates size categories associated with the four size bins detailed in the legend and D).

The dominating losses in large colonies were not fully reflected in percent changes of colony sizes (Figure 2D). Even though most large colonies decreased, and some small colonies showed very large percent increases, the relationship between percent change and previous size, latitude or date of first detection did not explain any significant variation in the data (glm, p > 0.05, r^2^ = 0.198), suggesting that large colonies remained large and the skewed size distribution in the North-East Atlantic gannet metapopulation generally, remained similar. See supplementary Figures 5, 6 and 7 for the individual relationships between the explanatory variables and percent change.

### New colony foundations, colonisation attempts and extinctions

Since 2013/14 three new GB gannet colonies were established; two in Scotland (St. Abbs, Marwick Head), one in Wales (Middle Mouse), with an additional colonisation attempt recorded in Scotland (Duncansby Head, Figure 1). New colonies and colonisation attempts have also been noted in the wider North-East Atlantic, but only at the Northern fringe of the distribution (Figure 1). A second Russian colony appeared to have been established between 2013 and 2015 on Bol’shoi Anikeev Island (Ezhov & Krasnov 2024) and in Norway, seven colonisation attempts were recorded since 2020 (Borgvær, Frugga, Skjaaberget, Ravnholmen, Kuskjeret, Steinflesan and Mellom-Barden). Some of these sites had counts of more than 20 AOS and may qualify as colonies if a few more years of count data confirm persistence. Within the same period, two established Norwegian colonies (Store Ulvøyholmen and Store Forøya) went extinct, while a third previously extinct colony (Skittenskarvholmen) held gannets for two years before being abandoned again. These recent foundations and extinctions bring the total number of colonies in the North-East Atlantic metapopulation to 50.

### Long-term and large-scale colony growth patterns

The colonies with more complete count data time series indicate that numbers generally continued to increase after 2013 (Figure 3: e.g. Sule Skerry, Westray, Noss, Helgoland). Given similar expected continued population growth between 2013/14 and the outbreak of HPAI in 2022 throughout the metapopulations, the reported colony trends likely underestimate the true decline caused by HPAI. However, some colonies did show a stagnation of growth across their time series, even before the 2022 HPAI outbreak (e.g. Figure 3: the Channel Islands colonies, Sulnasker, Runde).

**Figure 3:**
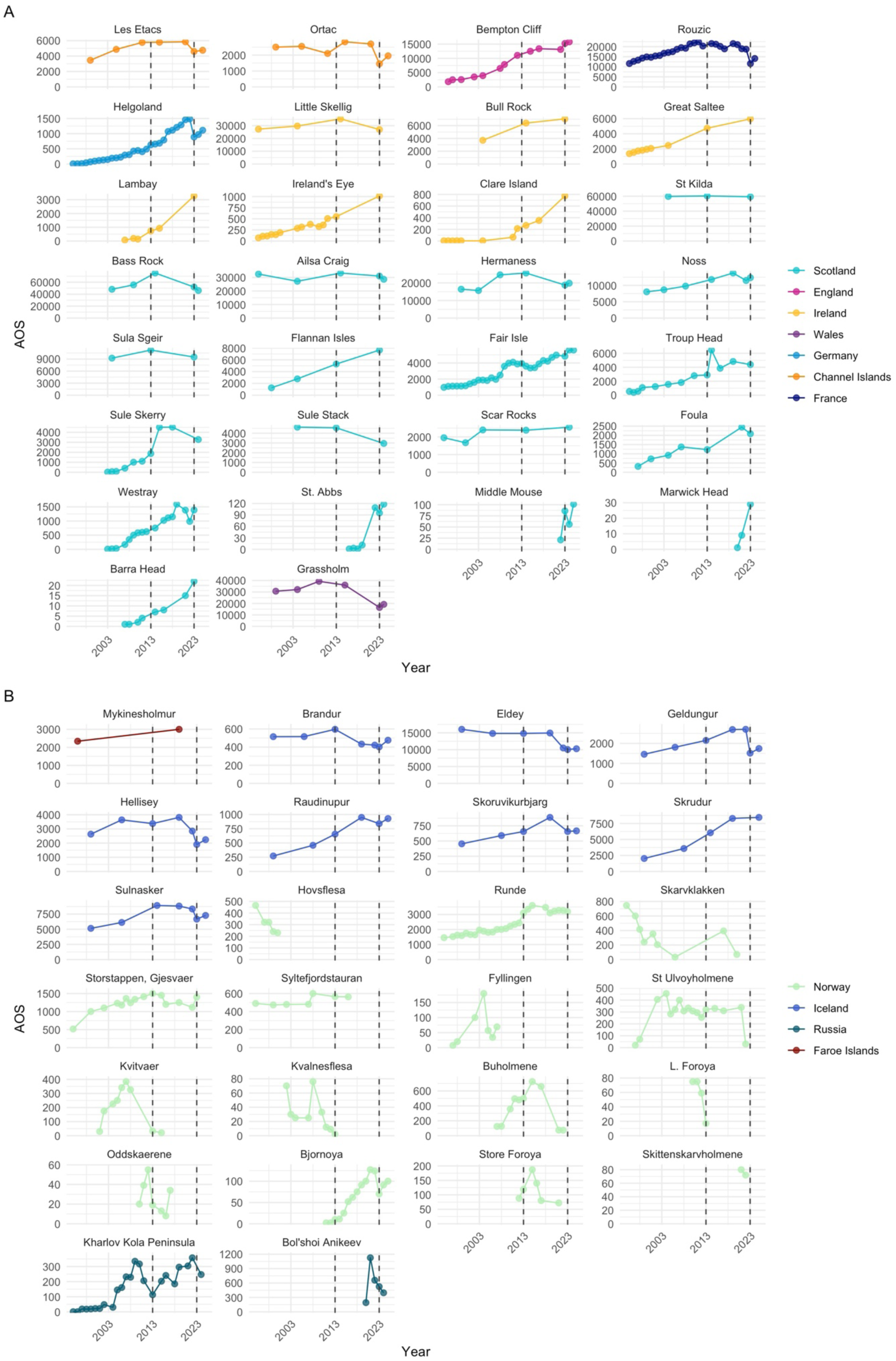
Colony trajectories for the A) central and southern part and B) northern part of the North-East Atlantic metapopulation since 1995. Dashed lines indicate the last two census windows 2013/14 and 2023/24 used here to calculate change in numbers. For longer-term colony trajectories since 1900 see Jeglinski *et al*. (2023). Note variable y axes.

### Detailed geographic accounts

#### Scotland

Scotland now contains 59% of the NE Atlantic population. This region holds 17 colonies, and 2 colonisation attempts (Table 3). The largest colony in Scotland (and in the North-East Atlantic) is once again St Kilda and no longer the Bass Rock (Murray *et al*. 2015, Nisbet *et al*. 2025), since the Bass Rock has seen the largest decline of all Scottish gannet colonies (31% decrease). St Kilda has a similar count to the 2003/04 census (Figure 3A). Some smaller colonies (less than 10, 000 AOS), such as Troup Head, Sule Skerry and West Westray, have seen some of the largest increases between the two censuses. Two new colonies - St Abb’s Head and Marwick Head - were founded since the last census. A colonisation attempt was recorded at Duncansby Head with a single nest in 2024. Even though there were previous records of birds breeding on Rockall (Wanless *et al*. 2015) and anecdotal evidence exists that in 2023 gannets were observed on nests from photos taken from a boat (L. Quinn *pers comms.*), no birds nested in 2024 when the aerial survey took place. Harvesting of chicks takes place at one colony, Sula Sgeir, where between 2004-2019 up to 2000 chicks were harvested each year (NatureScot licensing data). During 2020-2024 there was a voluntary suspension of the harvest from the harvesters, due to the Covid 19 and HPAI outbreaks.

**Table 3.** Summary of gannet colony counts in Scotland with comparisons across the most recent three gannet censuses, in order of size based on the 2023/24 census. Percentage changes are rounded to the nearest whole number. *marks colonisation attempts. n/a refers to those colonies that had not yet been established and therefore no % change given.

| Colony | 2003/04 (AOS) | 2013/14 (AOS) | % change between 2003-2013 | 2023/24 (AOS) | % change between 2013-2023 |
| --- | --- | --- | --- | --- | --- |
| St Kilda | 59,622 | 60,290 | +1 | 59,205 | -2 |
| Bass Rock | 48,065 | 75,259 | +57 | 51,844 | -31 |
| Ailsa Craig | 27,130 | 33,226 | +22 | 28,597 | -14 |
| Hermaness | 15,633 | 25,580 | +64 | 18,739 | -27 |
| Noss | 8,652 | 11,786 | +36 | 12,355 | +5 |
| Sula Sgeir | 9,225 | 11,230 | +22 | 10,200 | -9 |
| Flannan Isles | 2,760 | 5,280 | +91 | 7,646 | +45 |

| Colony | 2003/04 (AOS) | 2013/14 (AOS) | % change<br>between<br>2003-2013 | 2023/24 (AOS) | % change<br>between 2013-2023 |
| --- | --- | --- | --- | --- | --- |
| Fair Isle | 1,875 | 3,591 | +92 | 4,827 | +34 |
| Troup Head | 1,547 | 6,456 | +317 | 4,376 | -32 |
| Sule Skerry | 57 | 1,870 | +3,181 | 3,465 | +85 |
| Sule Stack | 4,618 | 4,550 | -1 | 2,947 | -35 |
| Scar Rocks | 2,394 | 2,375 | -1 | 2,551 | +7 |
| Foula | 919 | 1,226 | +33 | 2,086 | +70 |
| West Westray<br>(Noup) | 14 | 751 | +5,264 | 1,386 | +85 |
| St Abb's Head | n/a | n/a | n/a | 95 | n/a |
| Marwick Head | n/a | n/a | n/a | 29 | n/a - |
| Barra Head | n/a | 7 | n/a - | 22 | +214 |
| Duncansby Head* | n/a | n/a | n/a | 1 | n/a |
| Rockall* | n/a | 28 | n/a | 0 | -100 |
| **Scotland total** |  | 243,505 |  | **210,371** | **-14** |

#### Ireland

The island of Ireland holds six gannet colonies, comprising 13% of the NE Atlantic population, all in the Republic of Ireland. All colonies, except Little Skellig, increased since the 2013/14 survey (Table 4) (Newton *et al*. 2015), but a marked 24% reduction at Little Skellig (26,958 AOS, 60% of the total Irish population), resulted in an overall national population decline of 6%. A partial count of subsections of the Little Skellig colony from a 2018 drone survey indicated an 8.5% decrease between 2014 and 2018, suggesting that at least part of this population decline occurred prior to the 2022 HPAI outbreak (E. Murphy, *pers. comm*.; see also Figure 3). Increases at Bull Rock were modest as much of the ground surface and surrounding cliff faces are fully occupied by breeding pairs, limiting space for colony expansion. At Great Saltee, the 2023 surveys showed extensions of the colony to the east and west of the main colony and new sub-colonies on the southeastern and southwestern cliff faces. Lambay Island experienced the greatest percentage increase across all Irish colonies at 253%, including an extension of the colony to the east. Areas to the west of the main colony contained predominantly non-breeding / loafing individuals. Ireland’s Eye grew ∼200% since 2013/14, with nesting birds having spread to the island’s north facing cliffs. The small colony on Clare Island had some imagery overlap in the 2023 census, so the 2023 count may be a slight underestimate.

**Table 4.**
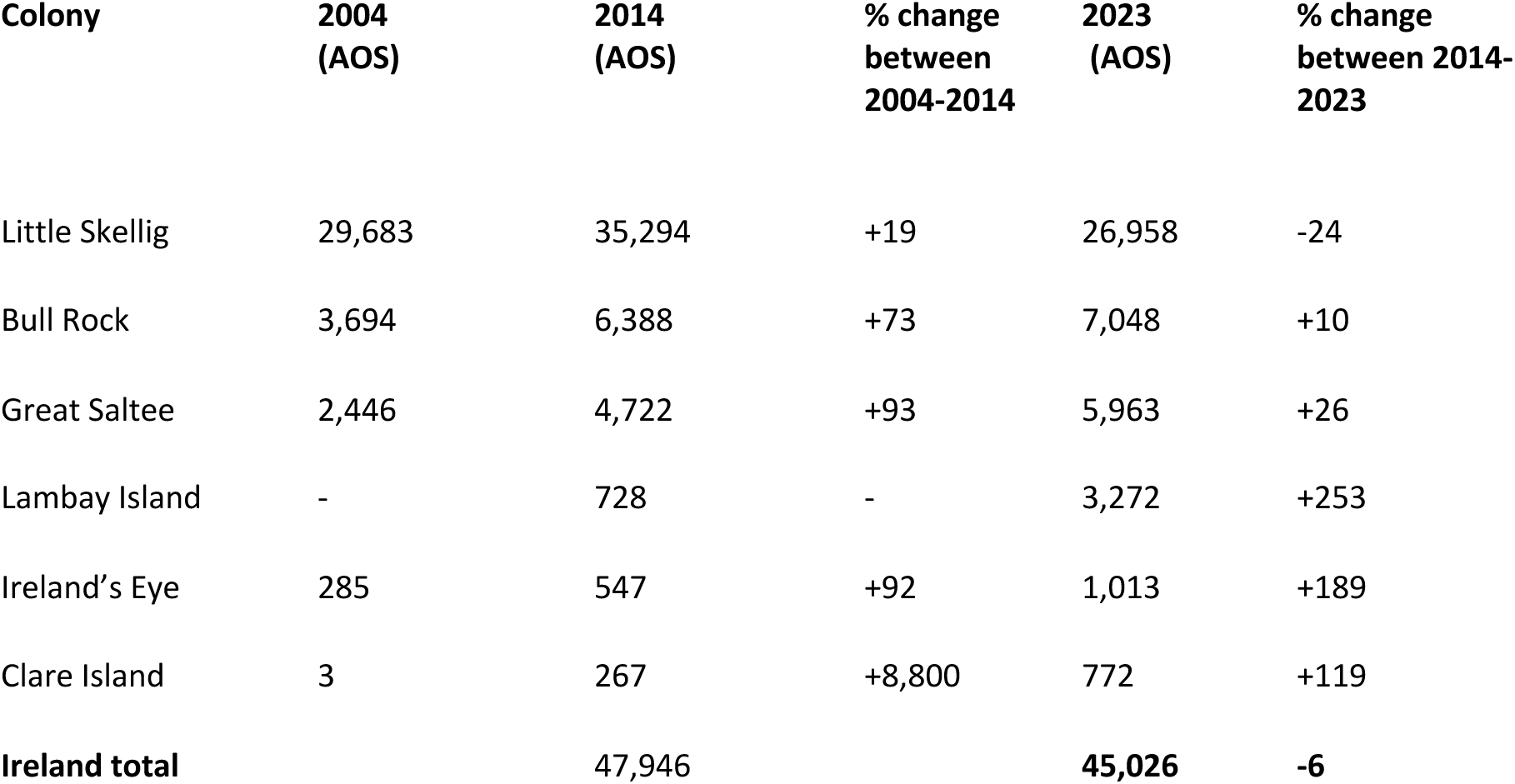
Summary of gannet colony counts in Ireland with comparisons across the most recent three gannet censuses. Percentage changes are rounded to the nearest whole number.

### Wales

Wales now contains 5% of the NE Atlantic population and holds two gannet colonies as of 2022 (Table 5). Grassholm, an island approximately 16 kilometres off Pembrokeshire in south-west Wales, is the main colony in Wales, whilst Ynys Padrig/Middle Mouse is a small new colony approximately 1 km off the north coast of Ynys Mon/Anglesey, North Wales. Gannets were first counted at Grassholm in 1872 with 12 birds recorded. The colony steadily increased up to 2009 when the largest estimated population was recorded at 39,292 AOS, and it became the third largest colony in the world (Figure 3A). In 2022, just before the HPAI outbreak, a colony count resulted in an estimate of 34,491 AOS. The 2023 count produced an estimate of 16,482 AOS, a 52% decline from the pre-HPAI 2022 count. The colony on Middle Mouse first started to establish in the summer of 2019, when more than a hundred gannets were noticed by a local boat operator. Breeders were noted in 2022 with 14 AOS present, which increased to 86 AOS in 2023.

**Table 5.**
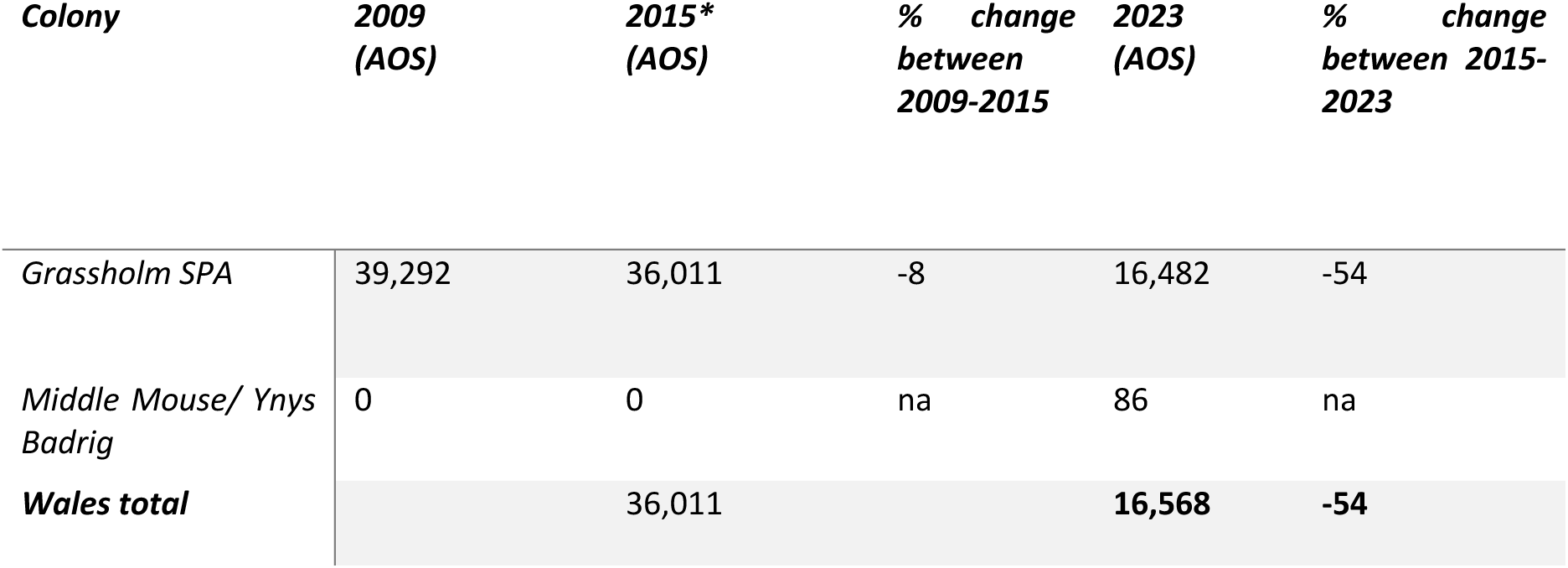
Summary of gannet colony counts in Wales with comparisons across the most recent three gannet censuses. Percentage change is rounded to the nearest whole number. *note that Murray *et al*. (2015) used the 2009 value for the 2013/14 census results. Here we instead compare between data collected in 2015, closer to the 2013/14 census time period, and the 2023 count.

### England

England now contains 4% of the NE Atlantic population. England holds a single colony at Bempton Cliffs within the Flamborough and Filey Coast SPA, founded in 1924. The colony grew by 73% between 2008 and 2012 surveys. However, growth declined to 38% between 2012 and 2023 (Table 6). The increase of 38% was unusual in the context of other large colonies over 10,000 AOS (Figure 2, panel C). A 2022 count, which was undertaken before the first impacts of HPAI in the colony were observed, was 13,125 AOS. Population trends differ between different sections of the colony. Staple Newk, previously the densest part of the colony, was significantly affected by HPAI and numbers here have declined, while other sections of the colony further north have increased (Butcher *et al*. 2023 & 2024).

**Table 6.** Summary of gannet colony counts in England with comparisons across the most recent three gannet censuses. Percentage change is rounded to the nearest whole number.

| *Colony* | *2008*<br>*(AOS)* | *2012*<br>*(AOS)* | *% change*<br>*between*<br>*2008-2012* | *2023*<br>*(AOS)* | *% change*<br>*between*<br>*2012-2023* |
| --- | --- | --- | --- | --- | --- |
| *Bempton Cliffs* | 6,392 | 11,061 | +73 | 15,233 | +38 |
| ***England total*** |  | 11,061 |  | **15,233** | **+38** |

### Channel Islands

The Channel Islands now contain 2% of the NE Atlantic population. The Channel Islands hold two gannet colonies, with a total of 6,036 AOS recorded across both colonies (Table 7). Les Etacs and Ortac both experienced declines in their populations since the last census, and it is notable that the Ortac colony was already in decline between the previous two censuses (Figure 3). Counts available from both 2015 (5,960 AOS Les Etacs, 2777 AOS Ortac) and 2021 (5,842 AOS Les Etacs, 2697 AOS Ortac), indicated that population growth at both colonies appears to have stagnated before HPAI (Figure 3), potentially due to the colonies reaching carrying capacity or low productivity (A. Purdie, pers. comm.).

**Table 7.** Summary of gannet colony counts in the Channel Islands with comparisons across the most recent three gannet censuses. Percentage changes are rounded to the nearest whole number.

| *Colony* | *2005*<br>*(AOS)* | *2011*<br>*(AOS)* | *% change*<br>*between*<br>*2005-2011* | *2023*<br>*(AOS)* | *% change*<br>*between*<br>*2011-2023* |
| --- | --- | --- | --- | --- | --- |
| *Les Etacs* | 4,862 | 5,765 | **+19** | 4,585 | **-20** |
| *Ortac* | 2,547 | 2,120 | **-17** | 1,451 | **-32** |
| ***Channel Islands***<br>***total*** |  | 7,885 |  | **6,036** | **-23** |

### Iceland

Iceland now contains 9% of the NE Atlantic population. Iceland holds eight long-established colonies, and a total of 30,484 AOS. In 2022, HPAI outbreaks in gannets appeared first in the south-western Icelandic colonies Eldey and the Westmann Islands group in April before spreading throughout the network of colonies globally (Lane *et al*. 2024), suggesting that the virus may have been introduced into the gannet metapopulation at these Icelandic breeding colonies. The south-western colonies (Eldey, Hellisey, Brandur, Geldungur, Sulnasker) all show marked decreases, while two of the three eastern colonies (Skrúður, Rauðinúpur) increased since the last census (Gardarsson 2019). Chick harvesting has taken place in Iceland although the numbers have been low in recent years (thought to be under 250 chicks annually since 2009) (Nature Conservation Agency of Iceland, 2026). Specific colonies are not reported in the hunting numbers but currently the majority comes from the eastern part of the country where the colonies Rauðinúpur, Skoruvík and Skrúður are situated. Harvesting from the south (including Westman Islands) and the west (Eldey) has been few chicks in the past 10 years.

**Table 8.** Summary of gannet colony counts in Iceland with comparisons across the most recent three gannet censuses, in order of size based on the most recent counts. Percentage changes are rounded to the nearest whole number. *colony count only available for 2025.

| *Colony* | *1999-2006*<br>*(AOS)* | *2013 (AOS)* | *% change*<br>*between 1999-*<br>*2013* | *2023 (AOS)* | *% change*<br>*between 2013-*<br>*2023* |
| --- | --- | --- | --- | --- | --- |
| Eldey | 14,812 | 14,810 | 0 | 10,049 | -32 |
| Sulnasker | 5,148 | 8,925 | +73 | 6,673 | -25 |
| Skrúður | 2,030 | 6,051 | +198 | 8,451* | +40 |
| Hellisey | 3,637 | 3,374 | -7 | 1,901 | -44 |
| Geldungur | 1,807 | 2,149 | +19 | 1,512 | -30 |
| Rauðínúpur | 272 | 655 | +141 | 839 | +28 |
| Skoruvíkurbjarg | 454 | 656 | +45 | 657 | 0 |
| Brandur | 516 | 596 | +16 | 402 | -33 |
| **Iceland total** |  | **37,216** |  | **30,484** | **-18** |

### France

France now contains 3% of the NE Atlantic population. France holds a single medium-sized colony Rouzic in the Sept-Ile marine reserve in Brittany, with a total of 11,592 AOS in the 2023 census. In 2023 the reserve perimeter expanded substantially from 280 to 19,700 hecatres, with a 130 hectare area closed to navigation from April 1^st^ to August 31^st^, incorporating the area where gannets reside (P. Provost, *pers comm.*). The colony has already been described as decreasing prior to the HPAI outbreak (Jeglinski *et al*. 2024, Grémillet *et al*. 2020) with severely reduced adult survival possibly due to competition with industrial fisheries (Le Bot *et al*. 2019) but also increasingly unsuitable environmental conditions modifying the breeding environment at the southern fringe of the gannet distribution (Jeglinski *et al*. 2024). A decrease of 46% AOS was noted between 2014 and 2023.

**Table 9.** Summary of gannet colony counts in France with comparisons across the most recent three gannet censuses. Percentage changes are rounded to the nearest whole number.

| *Colony* | *2003 (AOS)* | *2014 (AOS)* | *% change*<br>*between 2003-*<br>*2014* | *2023 (AOS)* | *% change*<br>*between 2013-*<br>*2023* |
| --- | --- | --- | --- | --- | --- |
| *Rouzic* | 16,718 | 21,545 | +29 | 11,592 | -46 |
| ***France total*** |  | **21,545** |  | **11,592** | **-46** |

### Norway

Norway now contains 2% of the North-East Atlantic metapopulation. Norway holds eight extant colonies, including Bjørnøya, the most northerly gannet colony globally, and a total of 5,799 AOS (Table 10). Skarvklakken appeared to be re-colonised in 2017 after having gone extinct in 2007 – the first occurrence of a re-colonisation of a previously abandoned site in the global gannet metapopulation. Norwegian colonies are generally characterised by distinct, and for the species, unusually unstable dynamics, with multiple colonisation/extinction events (Figure 3B). The gannet metapopulation expanded into the region in mid-1940s and expanded further north in the 1960s (Barrett 2008). The region has held a total of 15 colonies (Figure 3B), and colonies Store Forøya and Store Ulvøyholmen, both in the Vesterålen region, went extinct in the time period described here. In addition, five colonisation attempts have been noted in the most recent census, but it is unclear if these were breeding pairs or aggregations of prospecting non-breeders.

**Table 10.** Summary of gannet colony counts in Norway with comparisons across the most recent three gannet censuses, in order of size based on the most recent counts. Percentage changes are rounded to the nearest whole number. *marks colonisation attempts. n/a refers to those colonies that had not yet been established and therefore no % change given.

| Colony | 2002-2007<br>(AOS) | 2013 (AOS) | % change<br>between 2002-2013 | 2016-2023<br>(AOS) | % change<br>between 2013-2023 |
| --- | --- | --- | --- | --- | --- |
| Bjørnøya | n/a | 10 | n/a | 70 | 600 |
| Buholmene | 124 | 498 | +302 | 73 | -85 |
| Oddskjærene | n/a | 19 | n/a | 34 | +79 |
| Runde | 1,960 | 3,098 |  | 3,189 | 3 |
| Skarvklakken | 0 | 0 | n/a | 70 | n/a |
| St Ulvøyholmen | 455 | 319 | +58 | 0 | -100 |
| Store Forøya | n/a | 119 | n/a | 0 | -100 |
| Storstappen,<br>Gjæsvær | 1,100 | 1,500 | +36 | 1,400 | -7 |
| Syltefjordstauran | 479 | 565 | +18 | 563 | 0 |
| Borgvær* | n/a | n/a | n/a | 2 | n/a |
| Frugga* | n/a | n/a | n/a | 12 | n/a |
| Kuskjeret* | n/a | n/a | n/a | 64 | n/a |
| Ravnholmen,<br>Auvær* | n/a | n/a | n/a | 97 | n/a |
| Skjåberget* | n/a | n/a | n/a | 200 | n/a |
| Steinsflesan* | n/a | n/a | n/a | 25 | n/a |
| **Norway total** |  | **6,128** |  | **5,799** | **-5** |

### Faroe Islands

The Faroe Islands contain 1% of the NE Atlantic population. The Faroe Islands hold one long-established gannet colony, Mykineshólmur on the island of Mykines, the westernmost island in the Faroe archipelago. Counts have been infrequent and the most recent estimate, in 2023, provided a count of 3204 AOS, an increase from the previous count in 1996 (Table 11). As no gannet counts are available for the 2013/14 time period, unlike in other regions, only the 1996 and 2023 counts are presented in Table 11. The colony is harvested annually, taking an average of 495 gugas (fledgling chicks) in early September each year (data from 1989-2017, Olsen & Danielsen, unpublished).

**Table 11.** Summary of gannet colony counts in Faroes with comparisons across the most recent gannet counts for the Faroe Islands – including the census in 1996 and an estimate made in 2019. Percentage changes are rounded to the nearest whole number.

| Colony | 1996 (AOS) | 2023 (AOS) | % change<br>between 1996-2023 |
| --- | --- | --- | --- |
| Mykinesholmur | 2,340 | 3,000 | +37 |
| **Faroes total** | **2,340** | **3,204** | **+37** |

### Germany

Germany now contains <0.5% of the NE Atlantic population. Germany holds one colony on the island of Helgoland in the North Sea. The colony was established in 1991 and has subsequently been counted annually (Dierschke *et al*. 2023). This time series allows a more nuanced and detailed picture of the impact and recovery dynamics of the colony: after a period of rapid increase peaking in 2022 (pre-HPAI) with 1,485 AOS, numbers dropped drastically with a 40.2 % decrease between 2022 and 2023 followed by incipient recovery (Figure 3A). This highlights that the percent increase between 2013 and 2023 census is not illustrative of the impact of HPAI on the colony.

**Table 12.** Summary of gannet colony counts in Germany with comparisons across the most recent three gannet censuses. Percentage changes are rounded to the nearest whole number.

| Colony | 2003 (AOS) | 2014 (AOS) | % change<br>between 2003-2014 | 2023 (AOS) | % change<br>between 2014-2023 |
| --- | --- | --- | --- | --- | --- |
| Helgoland | 145 | 656 | +352 | 887 | +35 |
| **Germany total** |  | **656** |  | **887** | **+35** |

### Russia

Russia contains <0.5% of the NE Atlantic population. Russia now holds two colonies: the Kharlov Kola colony, established in the 1990s (Krasnov & Barrett 1997) and the newly established colony at Bol’shoi Anikeev, where gannets were first recorded breeding in 2015 and which held a total of 644 AOS in 2023. The newer colony has been established between the nearest Norwegian colony Syltefjord and the Kharlov Kola colony in Russia, thus does not represent a geographic breeding range expansion of the species towards the east.

**Table 13.** Summary of gannet colony counts in Russia with comparisons across the most recent two gannet censuses, in order of size based on the most recent counts. Percentage changes are rounded to the nearest whole number.

| *Colony* | *2004 (AOS)* | *2014 (AOS)* | *% change*<br>*between 2004-*<br>*2014* | *2023 (AOS)* | *% change*<br>*between 2014-*<br>*2023* |
| --- | --- | --- | --- | --- | --- |
| *Kharlov Kola Peninsula* | 30 | 113 | +277 | 247 | +119 |
| *Bol'shoi Anikeev* | n/a | 192 | n/a | 397 | +107 |
| ***Russia total*** |  | 305 |  | **644** | **+111** |

## Discussion

Our aim was to collate the most recent gannet colony counts for the entire North-East Atlantic metapopulation to understand changes in colony sizes following the severe HPAI outbreak in 2022. We document an overall 17% decrease across the North-East Atlantic metapopulation between 2013/14 and 2023/24, with nine of the 12 largest colonies (those over 10,000 AOS) experiencing a decline. This is a reversal of the previously documented continuous population increase and the first time a decline has been seen since gannets were protected from persecution in the late 19th century (Nelson 2002). However, not all geographic regions followed this general trend: numbers in Russia, Germany, the Faroe Islands and England increased while total numbers in all other regions decreased (though colonies within regions varied in increases or decreases). The rates of increase and decrease varied markedly between regions, with the highest proportional increases in Russia, at the northern extent of the distribution, and the highest proportional decrease in Wales and France, at the south-western fringe of the distribution.

### Drivers of metapopulation change

Mortality associated with HPAI was the likely driver of this large, sudden and unprecedented decrease across much of the species’ range (Lane *et al*. 2023, Giralt-Paradell *et al*. 2023; Atkinson *et al*. 2025). In 2022, HPAI outbreaks likely affected the species across its global distribution, with excessive mortality associated with HPAI outbreaks documented at 40 of the 53 extant colonies, with a lack of information for the remaining ones and only one confirmed case of absence of HPAI at a colony (Lane *et al*. 2024). As a long-lived species with generally very high adult survival rates (Wanless *et al*. 2006; Deakin *et al*. 2019), mortality of breeding adults has disproportionate effects on gannet population sizes, and influences the speed at which such populations can recover (Schreiber & Burger 2001). Efforts are underway to estimate HPAI-induced mortality of the adult breeding component (Matthiopoulos *et al*. in review; Lane *et al*. in review). The HPAI virus is constantly mutating and evolving, and the risk that future strains may pose is unknown. Furthermore, our understanding of immune responses to the virus in seabirds is in its infancy, as is our understanding of sublethal effects and likely timescales over which populations might recover.

In gannets, survival of the HPAI outbreak and HPAI antibodies have been shown to be related to changes in eye-colour from the normal pale blue irises to black or flecked irises (Lane *et al*. 2024). These changes are obvious to the naked eye and were noted at several colonies in and after 2022. In 2023, at Bass Rock and Bempton Cliffs, 9.6% and 9.4% of monitored birds were recorded with black eyes, respectively (Lewis *et al*. 2025). Black eyes were recorded in between 13 and 15 % of gannets in Russia (Ezhov & Krasnov 2024), and in around 30% of gannets in the Channel Islands in 2023 (Purdie, 2024), but we caution that these observations are generally collected opportunistically or from observation plots (i.e. not a complete colony survey), and methodological differences in recording complicate a direct comparison between colonies and conclusions on colony-specific survival rates. However, the geographically wide-spread observations of this phenotypic change support the notion of a ubiquitous and severe impact of HPAI on the global gannet population and that associated decreases in colony and metapopulation size documented here are largely attributable to it. Black-eyed gannets did not differ in breeding success (Lewis *et al*. 2025) or in their foraging behaviour (Ponchon *et al*. 2026) from that of gannets with normal eye colour, although understanding the potential sub-lethal effects of previous virus infection is still in its infancy. It should be noted that eye colour changes are no absolute signal of previous HPAI infection and that further quantitative research into this phenomenon is required.

HPAI did not only impact adult breeder survival but also impacted productivity and/or chick survival. On the Bass Rock, nest sites in a study area decreased by 75 % during the outbreak in 2022, and an index of productivity was estimated at 0.247 chicks per nest (Lane *et al*. 2024). Monitoring in the years following the outbreak at Grassholm, Bass Rock and Bempton, suggested that breeding success in these years was lower than the long-term average: at Grassholm productivity was 0.26 chicks per nest in 2024 compared to an average of 0.59 (2015-2019) (Morgan & Stevens 2024), on the Bass Rock productivity in 2023 was 0.56 compared to a long-term average of 0.78, and at Bempton, productivity in 2023 was 0.62 compared to 0.81 (mean between 2009 and 2021, Lewis *et al*. 2025 and references therein). HPAI thus impacted chicks directly in 2022 and potentially in subsequent years (though no observations of increased chick mortality beyond 2022 have been noted in the gannet), but also led to lower productivity in following years where no outbreaks happened. These reductions may relate to multiple factors. For example, reduced nest density at colonies as a direct consequence of high breeder mortality and slow infilling of gaps may facilitate predation on eggs by gulls. Also, large patches of vacant nest sites noted at e.g. Bass Rock (Harris *et al*. 2023) may have allowed recruitment of more first-time breeders, with limited experience (Morgan & Stevens 2024; Lewis *et al*. 2025) – these have been shown to have lower breeding success than established breeders (Nelson & Nelson 1963). Reductions in productivity may reduce the number of future recruits which will only become apparent in population trajectories with a lag of about five years, due to the delayed maturity of the gannet (Nelson 2002), suggesting that population recovery of the gannet may be slow. Whilst consideration of productivity data was not the focus of this current paper, these select examples emphasise the importance of collecting other demographic data beyond abundance (O’Hanlon *et al*. 2024).

Whilst the changes documented here are likely predominantly attributable to the HPAI outbreak in 2022, we note that general colony growth dynamics likely also played an important role, particularly in explaining colony-specific differences. Previous work has estimated an annual growth rate of ∼ 2 % comparing data from Operation Seafarer in 1969-70 to Seabird Counts data in 2013-2021 for large and long-established colonies (Wanless *et al*. 2023). Such high and continuous growth is likely only maintained in a colony not yet constrained by density dependence, where competition for space or resources is expected to reduce growth rates as colonies reach their carrying capacity leading to plateauing colony trajectories (Lewis *et al*. 2001). Indeed, there were indications that some colonies, such as Rouzic (Grémillet *et al*. 2020) and Little Skellig were already declining prior to the outbreak while other colonies had plateaued (e.g. the Channel Islands, Sule Sgeir, Ailsa Craig) (see Figure 3 A & B), or at least fallen short of the expected growth rate (e.g. all Irish colonies, except one), despite an overall increase in the North-East gannet metapopulation prior to 2022. A recent study on the impact of climate on gannet metapopulation dynamics illustrated climate-driven colony-specific differences in growth dynamics forecasting the decline of 15% of colonies in the near future, while 46% of colonies were forecast to plateau around their respective carrying capacity and, for an unmitigated climate change scenario, the overall metapopulation to plateau in ∼ 30 years (Jeglinski *et al*. 2024). Such colony-specific differences in density dependent constraints form the backdrop against which the HPAI outbreak played out, likely contributing to the differentiated colony-specific impacts that we observe in the colony count data.

### Spatial variation in change

Trends of colony size changes were not uniform, across the metapopulation or within regions, but varied both in direction and strength (Figure 2). We explored several plausible drivers of such variation: we hypothesized that larger colonies in which gannets may breed at higher densities may have experienced greater mortality due to increased transmission of HPAI and thus higher percent decreases. We also explored the date of detection of the first HPAI-related mortality in 2022, indicative of the spatio-temporal progression of the spread of the disease through the network of colonies. Finally, we explored the effect of latitude, which may affect HPAI outbreak dynamics as well as variation in suitability due to environmental conditions across the 26° latitude difference covered by the colony network. We showed that larger colonies lost more birds than small colonies and thus that the bulk of the reduction in numbers originated in the loss of tens of thousands of AOS in most of the largest colonies. However, these reductions were not proportional to colony size, i.e. larger colonies did not experience more severe percent reductions in numbers than smaller ones, negating our hypothesis that large colonies were more severely affected than small ones. The latter finding highlights that the heavily biased distribution of AOS between gannet colonies, towards few very large and many small colonies, was maintained after the HPAI outbreak.

We did not find support for a role of geographic distribution of colonies or the temporal pattern of HPAI spread though the gannet colony network in explaining variation in colony size changes, however there are other plausible differences between colonies that may contribute to the documented differences. Anecdotally, fieldworkers present in the colonies during the HPAI outbreak noticed variation in mortality between flatter parts of colonies and steeper cliffs (Jeglinski & Lane pers.obs; B. Olsen pers.obs; Nisbet *et al*. 2025). Topographic differences within and between colonies may contribute to differences in contact rates between neighbouring gannets due to differing densities, and to differences in the accumulation of corpses that may represent a source of infection. Topography may thus influence HPAI transmission dynamics and resulting mortality. This hypothesis could be explored in future research using the fine-scale high resolution 3D colony models that are generated as part of the image data processing of drone surveys. Quantifying these colony-specific topographical characteristics may better our understanding of colony-specific vulnerability to future disease outbreaks.

### Additional pressures on a reduced metapopulation

The reduction of the breeding gannet population presented in this census highlights the vulnerability of this species and of seabird populations in general (e.g. see also Tremlett *et al*. 2024; Indykiewicz *et al*. 2025) to the novel threat of HPAI. The severely reduced colony sizes documented here also urgently highlight the importance of reducing ongoing pressures that have the potential to detrimentally impact the species at the population-level, such as overfishing, bycatch, offshore renewables, and climate change, in order to increase the resilience of the gannet population. In addition to colony-specific variation in the strength of density-dependent growth regulation, acute stressors may also impact colonies differently, driving changes in colony trajectories over time in addition to HPAI mortality. For example, increasing predation pressure from reintroduced and increasing populations of white-tailed eagles *Haliaeetus albicilla* may have contributed to the unusual dynamics and extinction events in Norway (Barrett 2008). Mortality due to entanglement may also be negatively affecting populations. Some colonies may be particularly exposed to abandoned floating fishing gear (Bond *et al*. 2012), and present a considerable risk of entanglement both in the colony and where they are encountered and collected at sea. Several colonies during this census have observed either fishing materials incorporated into nests and/or lethal entanglement events (for example at Little Skellig, Ireland; Storstappan, Norway; the Channel Islands, Flannans, Sule Stack, Bempton, Helgoland, Runde and several Welsh colonies) (Personal observations from authors; Purdie 2024; Votier *et al*. 2011; Olaya Meza & Jones 2025). A standardised, repeatable protocol has been suggested to better monitor this wide-spread threat (O’Hanlon *et al*. 2017) and to improve assessment of population-level consequences. Scavenging on fishery discards plays an important role in the diet of breeding gannets (Bodey *et al*. 2014; Patrick *et al*. 2015; Giménez *et al*. 2021) and as such they are also commonly caught as bycatch in trawl fishing (Phillips *et al*. 2024), gillnets and longline fishing (Ramírez *et al*. 2024) in the North Atlantic. This occurs both during the breeding and non-breeding season. For example, longline fisheries alone catch an estimated minimum estimate of 18,500 gannets annually in the North-East Atlantic region (Ramírez *et al*. 2024). Data is once again lacking to be able to robustly quantify mortality (Araújo *et al*. 2022). Finally, harvesting of gannet fledglings occurs at one colony in Scotland, Sula Sgeir, as well as in the eastern and southern colonies in Iceland and in Mykines, Faroe Islands (Wanless *et al*. 2023; Jeglinski *et al*. 2023). Harvesting may have a direct impact on annual productivity (Wanless *et al*. 2023), can limit or modulate population growth, and may affect recruitment of local or external first-time breeders (Trinder, 2016). Depending on the level of harvest a net immigration from other nearby colonies will be required to sustain the population being harvested (Trinder, 2016), resulting in the harvested colonies potentially acting as population ‘sinks’. An up-to-date assessment of the impacts of harvesting on gannet populations across their range in the context of other pressures, as well as a better understanding of immigration/emigration dynamics at harvested colonies, would increase our understanding of the impact of harvesting chicks on the population growth rate of the colony itself and of neighbouring colonies.

### Advancement of survey techniques

Drones played an important role in the 2023/24 survey, the first time they have been used in a national gannet census. Such technological advancements can contribute towards improvement of censusing by increasing accessibility, standardisation, cost-efficiency and repeatability. When appropriate guidelines for reducing disturbance or injury to birds are adhered to, drones can provide clear images of colonies that would otherwise be difficult to reach by surveyors (Edney *et al*. 2023). In addition, these images can be kept and re-examined, compared to ‘one-off’ counts by human observers, with high-resolution images improving identification of AOS. Artificial intelligence was not used to count gannets for this census, although it is recognised that this is a rapidly developing field (e.g. Weinstein *et al*. 2022; Tyndall *et al*. 2024; Arnberg *et al*. 2025; Walker *et al*. 2025). In this census we did not require counters to record the number of loafing birds, however this could be considered in future surveys as it may be an indication of birds not breeding due to environmental or other conditions. In colonies where drones were used as the survey technique, no discernible disturbance response from gannets was observed (Harris *et al*. 2023; Nisbet *et al*. 2025). However, it is vital that drone pilots surveying seabird colonies continue to adhere to guidelines and provide details of any disturbance to wildlife within survey reports and/or publications. Within this current census, the use of aerial surveys also proved an efficient way to cover several colonies over a very short space of time (as per Scottish western colonies, Channel Islands, Ireland and Norway). As image resolution has improved over time, there is no longer a need for controlled disturbance flight to flush non-breeders, as was used previously (Murray *et al*. 2015).

### Data limitations

There are limitations in the data underlying the population changes we document here. Whilst the aforementioned technological advancements have aided the census, survey methods have changed at some of the colonies from fixed wing aerial or land/boat based to using drone technology for the first time. Comparisons of counts using more traditional methods alongside the newer drone technologies would be useful to ensure any changes in population reflect true change rather than a change in survey methodology. In addition to breeders, immature birds and adults on sabbatical may be present at colonies, which could potentially have led to an overestimate of AOS. The use of imagery may alleviate some of this error as counts can be checked retrospectively by multiple observers until consensus can be reached. Change between the two census periods is detected using only two points in time. Without more complete time series of colony counts, more complicated growth trajectories will be missed as well as the specific impact of HPAI between 2022 and 2023. This is highlighted in Figure 3 where we collated all available data for each colony, and where the few complete time series allowed us to trace the drop in gannet numbers that is likely directly associated with the impact of HPAI. We have shown that for Helgoland such detailed comparison reverses the direction of change, from 35 % increase between 2014 and 2023 to a reduction of 40% between 2022 and 2023 directly attributable to HPAI (Figure 3A). Similarly, the 2018 count for Sule Skerry was 4,515 AOS (Harris *et al*. 2019; Wanless *et al*. 2023), but in 2024 was 3,460 AOS. This represents a 30% decrease in the population between 2018 and 2024, even though the population has increased overall since the last national census in 2013/14. We therefore caution that the population changes presented here, despite being largely influenced by HPAI, do not accurately represent the full impact of HPAI on the numbers of adult breeding gannets in the metapopulation. Such impacts require estimates of HPAI-related mortality e.g. from mark recapture studies (Lane *et al*. in review) or models based on carcass observations (Matthiopoulos *et al*. in review). These are best estimated using appropriate models accounting for relevant factors that drive colony and metapopulation dynamics in the affected species to understand HPAI-related impacts, forecast colony recovery trajectories and highlight colonies and regions that are particularly sensitive to future disease outbreaks (Jeglinski *et al*. in prep.).

The changes we document here represent colony-specific dynamics and recovery, pressures and HPAI-related decreases (see above). In addition, we caution that numbers counted before and after the HPAI impact may not be directly comparable. HPAI mortality has created gaps in previously dense, regular breeding associations and new recruits sometimes appear to be infilling these gaps, rather than aggregating at fringes of dense associations. These birds may be loafers, immature birds prior to recruiting, or experienced birds that have lost their mates, which would be counted as AOS, but may not actually be contributing to the breeding population. While they may ultimately help colony recovery, the recruitment and pair bonding process can take several years in gannets and breeding success of new pairs may be lower than that of established ones (Nelson & Nelson 1963). Recruitment dynamics following the HPAI outbreak are currently poorly understood, but there is evidence for higher attendance of immature gannets (Sceviour *et al*. 2025), and non-breeding adults (Butcher *et al*. 2024), among the breeding pairs and observations of absence of traditional club sites at the fringe of colonies (Jeglinski pers. obs.). This suggests that the colony census here may be overestimating the current size of the breeding population, and therefore underestimating the impact of HPAI. It is vital to direct future research towards understanding the changed composition of colonies better to fill this knowledge gap.

### Conclusions

Gannet populations are vulnerable to several pressures summarised above, including disease (Lane *et al*. 2024), predation (Barrett *et al*. 2008), bycatch (Ramírez et al. 2024), entanglement (O’Hanlon *et al*. 2017), changes in food supply (Clark *et al*. 2020), harvesting (Trinder, 2016) pressures from offshore wind farm developments (Lane *et al*. 2020, Peschko *et al*. 2021) and climate change (Jeglinski *et al*. 2024). The updated colony counts we collate and discuss here represent the new ‘status quo’ for the North-East Atlantic metapopulation which forms the baseline for status assessments and conservation management decisions in the context of these multiple threats to the species. The decade-long gaps in national population monitoring for gannets across many of the North-East Atlantic colonies have likely masked significant population changes. Given the logistical challenges (even with drone-based technologies now available) and the cost of conducting regular national censuses, particularly at more remote colonies, a network of sentinel colonies that are surveyed at more frequent intervals could provide vital early warnings of population change between broader census periods. We are fortunate in that there are already sites that have regular time series data. The collection of demographic data beyond abundance, such as productivity and survival, for breeders but also other vulnerable age classes such as juveniles, would importantly benefit our understanding of population changes and the drivers of these changes in gannets. The importance of working together across nations has also been highlighted in this work, as only then do we get an understanding of large-scale variation in population responses across its entire North-East Atlantic metapopulation.

## Acknowledgements

None of this would be possible without the tremendous amount of work carried out by the fieldworkers who collected the data and funding bodies who have made it possible.

We thank the following individuals who were involved in the vital data collection, processing and/or analysis including: Sarah Harris (BTO). Scottish colonies: Connie Tremlett (RSPB); Bass Rock: Mike P Harris (UK Centre for Ecology & Hydrology), Sue Lewis (Edinburgh Napier University), Caroline J Nichol, Tom Wade, Amy Tyndall (University of Edinburgh) Maggie Sheddan (Scottish Seabird Centre). Foula: Joanne Monaghan, Rebecca Jackson, Andrew Stronach, Christian Carley, Jacob Thomas, Catherine Ramsay, Lucy Williamson, Miguel Hernández González. Hermaness: Mike Pennington; Noss: Jen Clark and Sally Reay (NatureScot). Western Scottish aerial surveys: Cristobal Olaya Meza, Ricki McCloud, Simon Warford, Georgia Catherine Anne Jones (APEM Ltd), Rachel Laurenson; St Kilda 2023: National Trust for Scotland: Alice Edney, Rachel Bonnici, Hebe Boyd-Wallis, Julie Watson, Mike Harris (UK Centre for Ecology and Hydrology); Noup Cliffs 2023: Matthew Marsh, Adam Martin, Alan Leitch, James Wilson, Lucy Mortlock, Chris Bell, Liz Fricker, Sophie Smith, Beth Mullier, Mike Pearce, Sandie Andrews. Marwick Head: Alan Leitch, Raymond Besant. Troup Head: Richard Humpidge, Edward Grace, Andrew Stronach, Allan Perkins. Sula Sgeir 2023: Liz Scott, Andy Scott, Derren Fox, Nicola Morley. Ailsa Craig 2023: Richard Humpidge, Derren Fox, Nicola Morley. Ailsa Craig and Scar Rocks 2024: Richard Humpidge, Sarah Dalrymple and Scarlett Humpidge. Welsh colonies: Grassholm: Greg Morgan, Nia Stephens and Richard Humpidge, Middle Mouse: Peter Evans and Jon Shaw. Irish colonies: Alyn Walsh, Andrew Power, Manon Clairbaux, Jamie Darby, Oriol Giralt Paradell. English colony: Bempton Cliffs 2023 and 2024: Dave O’Hara, Dave Aitken, Richard Humpidge, James Butcher, Keith Clarkson, Trevor Charlton. France: Armel Deniau. Faroes: Jens-Keld Jensen, Esbern and Hans Meinhard í Eyðanstovu, Sjúrður Hammer and Morten Frederiksen. And all the remaining North-East Atlantic colony fieldworker and data processors.

We also thank all the funding bodies involved in ensuring that this important census could be carried out, including: NatureScot, ScotWind developers of the East and North East plan areas, Scottish Government (via the Scotmer programme), The Crown Estate (via the OWEC Programme), Natural England, Natural Resources Wales, Irish National Parks and Wildlife Service, Research Ireland, Royal Society for the Protection of Birds (RSPB) (including Tim and Kim Allen), Norwegian fieldwork was part of the SEAPOP programme (www.seapop.no/en), financed by the Norwegian Ministry of Climate and Environment via the Norwegian Environment Agency, the Norwegian Ministry of Energy via the Norwegian Research Council (grant 192141), and Offshore Norge. Zoe Deakin (RSPB) facilitated part-funding of Icelandic aerial surveys. The Faroe Island surveys were funded by the Aage V. Jensen Charity Foundation, and the drone equipment by the Brødrene Hartmanns Fond. And all other funding bodies from the remaining colonies.

