## Supplementary Table S1 for "An updated population estimate for Northern gannets across their north-east Atlantic breeding range following the 2022 outbreak of High pathogenicity avian influenza"

Author’s affiliations

3. School of Biodiversity, One Health and Veterinary Medicine, University of Glasgow, UK

4. British Trust for Ornithology, The Nunnery, Thetford

5. Scottish Seabird Centre, North Berwick

6. Norwegian Institute for Nature Research NINA, Trondheim, Norway

7 Faeroe Marine Research Institute Havstovan, Tórshavn, Faroe Islands

8. Institut fuer Vogelforschung ‘Vogelwarte Helgoland‘, Helgoland, Germany

9. Murmansk Marine Biological Institute of the Russian Academy of Sciences, Russia

10. Natural Science Institute of Iceland, Gardabaer, Iceland

11. School of Biomedical Sciences, Oxford Brookes University, Headington Campus, Oxford, OX3 0BP

12. School of Biological, Earth & Environmental Sciences, University College Cork, Ireland

13. Natural England, Foss House, YO1 7PX

14. Ligue pour la Protection des Oiseaux, Réserve Naturelle Nationale des Sept-Iles, Pleumeur Bodou, France

15. Alderney Wildlife Trust, St. Anne, Alderney, Bailiwick of Guernsey

16. Sustainability Institute, University College Cork, Ireland

17. Natural Resources Wales, Maes y Ffynnon, Penrhosgarnedd, Bangor, Gwynedd, LL57 2DW

18. The National Trust for Scotland, Balnain House, Huntly Street Inverness IV3 5HR

19. BirdLife Møre og Romsdal, Sandshamn, Norway

20. Norwegian Polar Institute, Post box 6606 Stakkevollan, Tromsø, Norway

21. National Parks and Wildlife Service, Dublin, Ireland

22. RSPB Centre for Conservation Science, Etive House, Inverness, IV2 3BW, UK

23. UK Centre for Ecology & Hydrology, Bush Estate, Penicuik, UK, EH26 0QB

24. RSPB Ramsey Island, St Davids, Pembrokeshire, SA62 6PY

25. Kandalaksha State Nature Reserve, Russia.

**Table S1.** Summary of all the colonies censused including: survey method used, drone types employed (where relevant), as well as references to additional data sources where more in depth reports on the colony surveys exist (left blank if no further reporting exists). The Seabird Monitoring Programme Site identifier (SMP SiteID) is shown for the GB, Ireland and Channel Islands colonies (<https://app.bto.org/seabirds/public/index.jsp>).

| **County** | **Colony** | **SMP SiteID** | **Survey method** | **Date(s) of survey 2023** | **Date of survey**  **2024** | **Census unit** | **Additional data source/details** |
| --- | --- | --- | --- | --- | --- | --- | --- |
| **Scotland** | Bass Rock | 96965 | Drone  DJI Matrice 300 RTK | 27/06/2023 | 29/07/2024 | AOS | Harris *et al.* 2023;  Burton *et al.* 2024. |
| **Scotland** | St Kilda (Boreray and Stacs) | 88585 | Drone  DJI Matrice 30T, Mavic 3T, Mini 3 Pro | 3/06/2023  14-15/06/2023 | - | AOS | Nisbet *et al.* 2025 |
| **Scotland** | Ailsa Craig | 86638 | Drone DJI Mavic 3 Pro | 11/07/2023 | 12-13/07/2024 | AOS | Humpidge 2024 |
| **Scotland** | Hermaness | 89616 | Land, boat, and photos from land and sea | 01/06/2023 | - | AOS | Pennington, 2023. |
| **Scotland** | Noss | 98196 | Land, boat and photos from land and boat | 2/07/2023 | - | AON | Noss NNR report 2023 |
| **Scotland** | Sula Sgeir | 84590 | Aerial survey | 21/7/23 | 18/6/2024 | AOS | Olaya *et al.* 2024 |
| **Scotland** | Flannan Isles | 84608 | Aerial survey | - | 18/6/2024 | AOS | Olaya *et al.* 2024 |
| **Scotland** | Fair Isle | 84981 | Land | 14/06/2023 | 01/06/2024 | AON | Fair Isle Bird Observatory Report No. 75 (2023); Fair Isle Bird Observatory Report No. 76 (2024) |
| **Scotland** | Troup & Lion's Head | 100802 | Land and drone | 9-13/6/2023 | - | AON | - |
| **Scotland** | Sule Stack | 84624 | Aerial survey | - | 18/6/2024 | AOS | Olaya *et al.* 2024 |
| **Scotland** | Sule Skerry | 98132 | Aerial survey | - | 18/6/2024 | AOS | Olaya *et al.* 2024 |
| **Scotland** | Foula | 98223 | Boat | 5/07/2023 | - | AOS |  |
| **Scotland** | Scar Rocks | 96547 | Drone DJI Mavic 3 Pro | - | 14/7/2024 | AOS | Humpidge 2024 |
| **Scotland** | Noup Cliffs RSPB (West Westray) | 84398 | Land | 12-15/6/23 | - | AON | Marsh *et al.* 2023 |
| **Scotland** | Rockall | 110480 | Aerial survey | - | 10/7/2024 | AOS | Olaya *et al.* 2024 |
| **Scotland** | St Abb's Head NNR | 85934 | Land and Boat | 03/06/2023 | 02/06/2024 | AON | Hatsell 2023; Hatsell 2024 |
| **Scotland** | Marwick Head | 90182 | Drone | 6/6/2023 | - | AON | - |
| **Scotland** | Berneray (Barra Head) | 84592 | Land | 2023 | - | AOS | CachiaZammit 2023:CachiaZammit 2024 |
| **Scotland** | Duncansby Head | 80522 | Land | - | 13/06/2024 | AON | SMP, 2025 |
| **Wales** | Grassholm | 81736 | Drone  DJI Mavic Pro | 20/6/2023 | 20/06/2024 | AOS | Morgan *et al.* 2023  Morgan & Stephens 2024. |
| **Wales** | Middle Mouse | 86189 | Boat | 10/08/2023 | 21/07/2024 | AOS |  |
| **Ireland** | Little Skellig | 97630 | Aerial survey | 29/06/2023 | - | AOS | Murphy *et al.* 2025 |
| **Ireland** | Bull Rock | 98272 | Aerial survey | 29/06/2023 | - | AOS | Murphy *et al.* 2025 |
| **Ireland** | Great Saltee | 97585 | Aerial survey | 29/06/2023 | - | AOS | Murphy *et al.* 2025 |
| **Ireland** | Lambay Island | 85335 | Aerial survey | 29/06/2023 | - | AOS | Murphy *et al.* 2025 |
| **Ireland** | Clare Island | 100780 | Aerial survey | 29/06/2023 | - | AOS | Murphy *et al.* 2025 |
| **Ireland** | Ireland's Eye | 85389 | Aerial survey | 29/06/2023 | - | AOS | Murphy *et al.* 2025 |
| **England** | Flamborough Head and Bempton Cliffs | 98273 | Boat survey | 16/06/2023 | 17/6/2024 | AOS | Butcher *et al.* 2023; 2024 |
| **Channel Islands** | Les Etacs | 98275 | Aerial survey | 04/07/2023 | - | AOS | Purdy, 2024 |
| **Channel Islands** | Ortac | 98274 | Aerial survey | 04/07/2023 | - | AOS | Purdy, 2024 |
| **Iceland** | Brandur | n/a | Aerial survey | 10/06/2023 | - | AOS | - |
| **Iceland** | Eldey | n/a | Aerial survey | 10/06/2023 | - | AOS | - |
| **Iceland** | Geldungur | n/a | Aerial survey | 10/06/2023 | - | AOS | - |
| **Iceland** | Hellisey | n/a | Aerial survey | 10/06/2023 | - | AOS | - |
| **Iceland** | Skoruvikurbjarg | n/a | Aerial survey | 12/06/2023 | - | AOS | - |
| **Iceland** | Skrudur | n/a | Aerial survey | 04/07/2025 | - | AOS | - |
| **Iceland** | Sulnasker | n/a | Aerial survey | 10/06/2023 | - | AOS | - |
| **Iceland** | Raudinupur | n/a | Aerial survey | 12/06/2023 | - | AOS | - |
| **Norway** | Buholmene | n/a | Aerial survey | 2022 | - | AOS | - |
| **Norway** | Oddskaerene | n/a | Aerial survey | 2017 | - | AOS | - |
| **Norway** | Runde | n/a | Drone DJI Mavic 3 Pro and Enterprise | 29/07/2023 | 03/08/2024 | AOS | - |
| **Norway** | Skarvklakken | n/a | Aerial survey | 2020 | - | AOS | - |
| **Norway** | St Ulvoyholmene | n/a | Aerial survey | 10/07/2023 | - | AOS | - |
| **Norway** | Store Foroya | n/a | Aerial survey | 10/07/2023 | - | AOS | - |
| **Norway** | Storstappen | n/a | Land | 2023 | - | AOS | - |
| **Norway** | Syltefjordstauran | n/a | Land | 2016 | - | AOS | - |
| **Norway (Svalbard)** | Bjornoya | n/a | Drone DJI Mavic 3 Pro | 29/07/2023 | 28/07/2024 | AON | - |
| **Norway** | Borgvaer | n/a | Aerial survey | 2023 | - | AOS | - |
| **Norway** | Frugga | n/a | Aerial survey | 2023 | - | AOS | - |
| **Norway** | Kuskjeret | n/a | Aerial survey | 2023 | - | AOS | - |
| **Norway** | Ravnholmen | n/a | Aerial survey | 2023 | - | AOS | - |
| **Norway** | Skjaaberget | n/a | Aerial survey | 2023 | - | AOS | - |
| **Norway** | Steinsflesan | n/a | Aerial survey | 2023 | - | AOS | - |
| **France** | Rouzic | n/a | Aerial survey | 12/05/2023 | 23/05/2023 (drone) | AOS | Provost *et al.* 2025; 2025 |
| **Germany** | Helgoland | n/a | Land count | 2023 | - | AOS | Dierschke *et al.* 2024 |
| **Faroe Islands** | Mykinesholmur | n/a | Drone DJI Matrice 300 RTK | 22-24/07/2023 | - | AOS | J.H.F. Castenschiold (pers comm.) |
| **Russia** | Bol’shoi Anikeev | n/a | Drone DJI Mavic 2 Pro | 10/06/2023 | 20/07/2024 | AON | Ezhov & Krasnov 2024 |
| **Russia** | Kharlov | n/a | Drone DJI Mavic 2 Pro | - | 21/07/2024 | AON | Ezhov & Krasnov 2024 |
