## Supplementary Table S2 for "An updated population estimate for Northern gannets across their north-east Atlantic breeding range following the 2022 outbreak of High pathogenicity avian influenza"

Author’s affiliations

3. School of Biodiversity, One Health and Veterinary Medicine, University of Glasgow, UK

4. British Trust for Ornithology, The Nunnery, Thetford

5. Scottish Seabird Centre, North Berwick

6. Norwegian Institute for Nature Research NINA, Trondheim, Norway

7 Faeroe Marine Research Institute Havstovan, Tórshavn, Faroe Islands

8. Institut fuer Vogelforschung ‘Vogelwarte Helgoland‘, Helgoland, Germany

9. Murmansk Marine Biological Institute of the Russian Academy of Sciences, Russia

10. Natural Science Institute of Iceland, Gardabaer, Iceland

11. School of Biomedical Sciences, Oxford Brookes University, Headington Campus, Oxford, OX3 0BP

12. School of Biological, Earth & Environmental Sciences, University College Cork, Ireland

13. Natural England, Foss House, YO1 7PX

14. Ligue pour la Protection des Oiseaux, Réserve Naturelle Nationale des Sept-Iles, Pleumeur Bodou, France

15. Alderney Wildlife Trust, St. Anne, Alderney, Bailiwick of Guernsey

16. Sustainability Institute, University College Cork, Ireland

17. Natural Resources Wales, Maes y Ffynnon, Penrhosgarnedd, Bangor, Gwynedd, LL57 2DW

18. The National Trust for Scotland, Balnain House, Huntly Street Inverness IV3 5HR

19. BirdLife Møre og Romsdal, Sandshamn, Norway

20. Norwegian Polar Institute, Post box 6606 Stakkevollan, Tromsø, Norway

21. National Parks and Wildlife Service, Dublin, Ireland

22. RSPB Centre for Conservation Science, Etive House, Inverness, IV2 3BW, UK

23. UK Centre for Ecology & Hydrology, Bush Estate, Penicuik, UK, EH26 0QB

24. RSPB Ramsey Island, St Davids, Pembrokeshire, SA62 6PY

25. Kandalaksha State Nature Reserve, Russia.

**Table S2.** Census results for all North-East Atlantic gannet colonies (AOS). Wherever possible 2023 was used as the count year, or 2024 if the 2024 count was more robust or there was no 2023 count available. The exceptions to using 2023 or 2024 related to no colony count taking place in 2023 or 2024. This resulted in once Icelandic, four Norwegian and one Faroe Islands colonies having a count either before or after 2023/24.

| **Country** | **Colony** | **Colony abbreviation** | **Counts used for updated census** | **Count unit** | **Year of count** |
| --- | --- | --- | --- | --- | --- |
| Scotland | St Kilda | SK | 59205 | AOS | 2023 |
| Scotland | Bass Rock | BsR | 51844 | AOS | 2023 |
| Scotland | Ailsa Craig | AC | 28597 | AOS | 2024 |
| Scotland | Hermaness | Hr | 18739 | AOS | 2023 |
| Scotland | Noss | Ns | 12355 | AON | 2023 |
| Scotland | Sula Sgeir | SlSg | 10200 | AOS | 2024 |
| Scotland | West Group (Flannans) | WG | 7646 | AOS | 2024 |
| Scotland | Fair Isle | FI | 4827 | AON | 2023 |
| Scotland | Troup & Lion's Head (Coast & Reserve) | TH | 4376 | AON | 2023 |
| Scotland | Sule Skerry | SlSt | 3465 | AOS | 2024 |
| Scotland | Sule Stack | SlSk | 2947 | AOS | 2024 |
| Scotland | Big Scar | BS | 2551 | AOS | 2024 |
| Scotland | Foula | Fl | 2086 | AOS | 2023 |
| Scotland | Noup Cliffs (West Westray) | Ws | 1386 | AON | 2023 |
| Scotland | St Abb's Head NNR | SA | 95 | AON | 2023 |
| Scotland | Marwick Head | MH | 29 | AON | 2023 |
| Scotland | Berneray (Barra Head) | BH | 22 | AOS | 2023 |
| Scotland | Duncansby Head | DH | 1 | AON | 2024 |
| Scotland | Rockall | n/a | 0 | AOS | 2024 |
| Ireland | Little Skellig - Whole Island | LS | 26958 | AOS | 2023 |
| Ireland | The Bull | BlR | 7048 | AOS | 2023 |
| Ireland | Great Saltee Island | GS | 5963 | AOS | 2023 |
| Ireland | Lambay | Lm | 3272 | AOS | 2023 |
| Ireland | Ireland's Eye | IE | 1013 | AOS | 2023 |
| Ireland | Clare Whole Island | CI | 772 | AOS | 2023 |
| Wales | Grassholm | Gr | 16482 | AOS | 2023 |
| Wales | Middle Mouse | MM | 96 | AOS | 2023 |
| England | Bempton Cliffs | Bm | 15233 | AOS | 2023 |
| Channel Islands | Les Etacs | LE | 4585 | AOS | 2023 |
| Channel Islands | Ortac | Or | 1451 | AOS | 2023 |
| Iceland | Eldey | El | 10049 | AOS | 2023 |
| Iceland | Skrudur | Skrd | 8451 | AOS | 2025 |
| Iceland | Sulnasker | Sl | 6673 | AOS | 2023 |
| Iceland | Hellisey | Hll | 1901 | AOS | 2023 |
| Iceland | Geldungur | Gl | 1512 | AOS | 2023 |
| Iceland | Raudinupur | Rd | 839 | AOS | 2023 |
| Iceland | Skoruvikurbjarg | Skrvkr | 657 | AOS | 2023 |
| Iceland | Brandur | Brn | 402 | AOS | 2023 |
| France | Rouzic | Rz | 11592 | AOS | 2023 |
| Norway | Runde | Rn | 3189 | AOS | 2023 |
| Norway | Storstappen, Gjesvær | Str | 1400 | AOS | 2023 |
| Norway | Syltefjordstauran | Sy | 563 | AOS | 2016 |
| Norway | Skjåberget | Skj | 200 | AOS | 2023 |
| Norway | Ravnholmen, Auvær | Rv | 97 | AOS | 2023 |
| Norway | Bjørnøya | Bj | 73 | AON | 2023 |
| Norway | Buholmene | Bh | 73 | AOS | 2022 |
| Norway | Skarvklakken | Skrvkl | 70 | AOS | 2020 |
| Norway | Kuskjeret | Ks | 64 | AOS | 2023 |
| Norway | Oddskærene | Od | 34 | AOS | 2017 |
| Norway | Steinsflesan | Stn | 25 | AOS | 2023 |
| Norway | Frugga | Fr | 12 | AOS | 2023 |
| Norway | Borgvær | Brg | 2 | AOS | 2023 |
| Norway | St Ulvøyholmene | SU | 0 | AOS | 2023 |
| Norway | Store Forøya | SF | 0 | AOS | 2023 |
| Faroe Islands | Mykinesholmur | My | 3204 | AOS | 2023 |
| Germany | Helgoland | Hlg | 887 | AOS | 2023 |
| Russia | Bol'shoi Anikeev | KKP | 397 | AON | 2024 |
| Russia | Kharlov Kola Peninsula | BA | 247 | AON | 2024 |
| **Total** |  |  | **345,854** | **AOS** |  |
